# Effects of age on the green autofluorescence of the skin and fingernails of healthy persons

**DOI:** 10.1101/449660

**Authors:** Mingchao Zhang, Yue Tao, Danhong Wu, Yujia Li, Xingdong Chen, Weihai Ying

**Affiliations:** Med-X Research Institute and School of Biomedical Engineering, Shanghai Jiao Tong University, Shanghai 200030, P.R. China; State Key Laboratory of Genetic Engineering and Collaborative Innovation Center for Genetics and Development, School of Life Sciences, Fudan University, Shanghai 200438, China; Fudan University Taizhou Institute of Health Sciences, Taizhou 225300, China; Human Phenome Institute, Fudan University, Shanghai 201203, China; Department of Neurology, Shanghai Fifth People’s Hospital, Fudan University, Shanghai, P.R. China

**Keywords:** Age, Autofluorescence, Skin, Fingernails, Age-independent diseases

## Abstract

Our recent studies have suggested that altered Pattern of Autofluorescence (AF) of skin and fingernails are novel biomarkers of multiple major diseases. The age of all of the subjects in these studies ranges from 50 to 80 years of old. For future studies on the potential diagnostic value of green AF for age-independent diseases, it is required to answer the following question: Are there differences in the green AF of healthy persons of various age populations? In our current study, we determined the green AF of the skin and fingernails of healthy persons in several age populations, showing significant age dependence of the AF: First, the green AF intensity of the age group between 15 to 20 years of old is significantly higher than that of the age group between 50 to 80 years of old at both right and left Centremetacarpus, right Dorsal Centremetacarpus, and right Dorsal Index Finger. Second, for the green AF intensity of the age groups of 15 to 20 years of old, 60 to 70 years of old, 70 to 80 years of old, and 81 to 85 years of old, the green AF intensity is negatively correlated with the age at both right and left Centremetacarpus and right Dorsal Centremetacarpus. Collectively, our study has provided first evidence indicating the age dependence of the AF intensity of human’s skin, which has established essential basis for the studies that determine the diagnostic value of the green AF for the age-independent diseases.

## Introduction

Our recent studies have suggested that characteristic ‘Pattern of Autofluorescence (AF)’ could be a novel biomarker for non-invasive diagnosis of acute ischemic stroke (AIS) (3), myocardial infarction (MI) (17), stable coronary artery disease (17), Parkinson’s disease (4) and lung cancer (10). In particular, there are marked and characteristic increases in the green AF intensity in the fingernails and certain regions of the skin of the AIS patients (3) and MI patients (17). Since oxidative stress can lead to increased epidermal green AF by inducing proteolytic cleavage of keratin 1 (8,9), the increased oxidative stress in the blood of AIS patients (2,6,13) and MI patients (1,5,7,11,16) may be responsible for the increased AF intensity.

The age of all of the subjects in these studies ranges from 50 – 80 years of old. For future studies on the potential diagnostic value of green AF for age-independent diseases, it is required to answer the following question: Are there differences in the green AF of healthy persons of various age populations? In our current study, we determined the green AF of the skin and fingernails of healthy persons in several age populations, showing significant age dependence of the AF.

## Methods and materials

### Human subjects

The healthy human subjects lived in the various districts of Shanghai, P.R. China. The human subjects in our study were divided into four groups: Group 1: The Age Group between 15 – 20 years of old; Group 2: The Age Group between 50 – 60 years of old; Group 3: The Age Group between 60 – 70 years of old; Group 4: The Age Group between 70 – 80 years of old; and Group 5: The Age Group between 81 – 85 years of old.

### Determinations of the Autofluorescence of Skin and Fingernails

A portable AF imaging equipment was used to detect the AF of the fingernails and certain regions of the skin of the human subjects. The excitation wavelength is 485 nm, and the emission wavelength is 500 - 550 nm. For all of the human subjects, the AF intensity in the following seven regions on both hands, i.e., fourteen regions in total, was determined, including the Index Fingernails, Ventroforefingers, Dorsal Index Finger, Centremetacarpus, Dorsal Centremetacarpus, Ventribrachium and Dorsal Antebrachium.

### Statistical analyses

All data are presented as mean ± SEM. Data were assessed by Student t test, except where noted. *P* values less than 0.05 were considered statistically significant.

## Results

### 1. The green AF intensity of the age group between 15 – 20 years of old is significantly higher than that of the age group between 50 – 80 years of old at certain regions examined

We determined the green AF intensity of the age group between 15 – 20 years of old and that of the age group between 50 – 80 years of old. We found that the green AF intensity of the age group between 15 – 20 years of old is significantly higher than that of the age group between 50 – 80 years of old at both right and left Centremetacarpus (Fig. 1), right Dorsal Centremetacarpus (Fig. 2), and right Dorsal Index Finger (Fig. 3). In contrast, no difference of green AF intensity between these two age groups at other regions examined, including Ventroforefinger (Supplemental Fig. 1), Ventriantebrachium (Supplemental Fig. 2), Dorsal Antebrachium (Supplemental Fig. 3), and Index Fingernails (Supplemental Fig. 4).

**Fig. 1.**
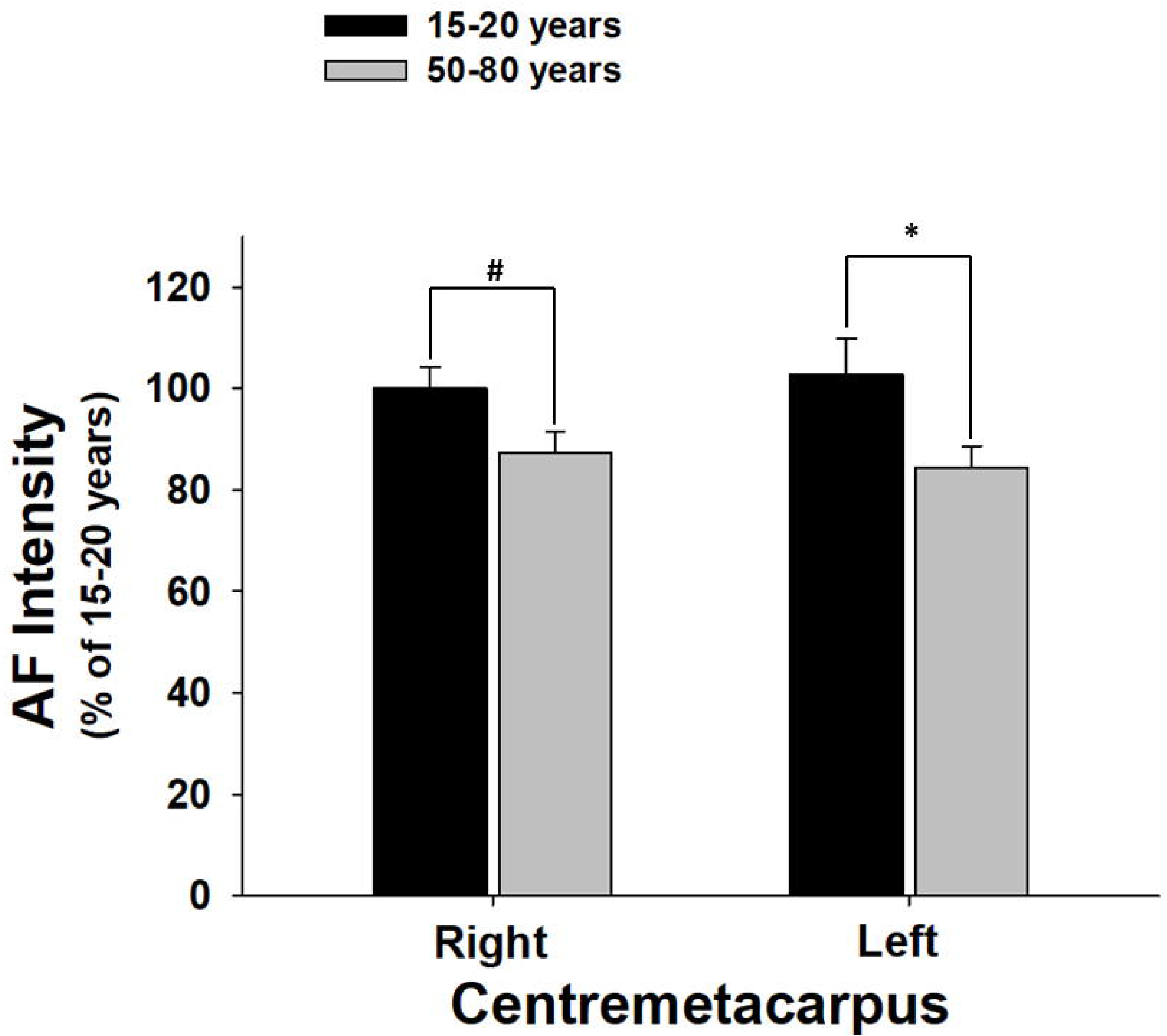
The green AF intensity of the age group between 15 – 20 years of old is significantly higher than that of the age group between 50 – 80 years of old at both right and left Centremetacarpus. The number of subjects in the age group between 15 – 20 years of old and the age group between 50 – 80 years of old is 50 and 73, respectively. *, *p* < 0.05.

**Fig. 2.**
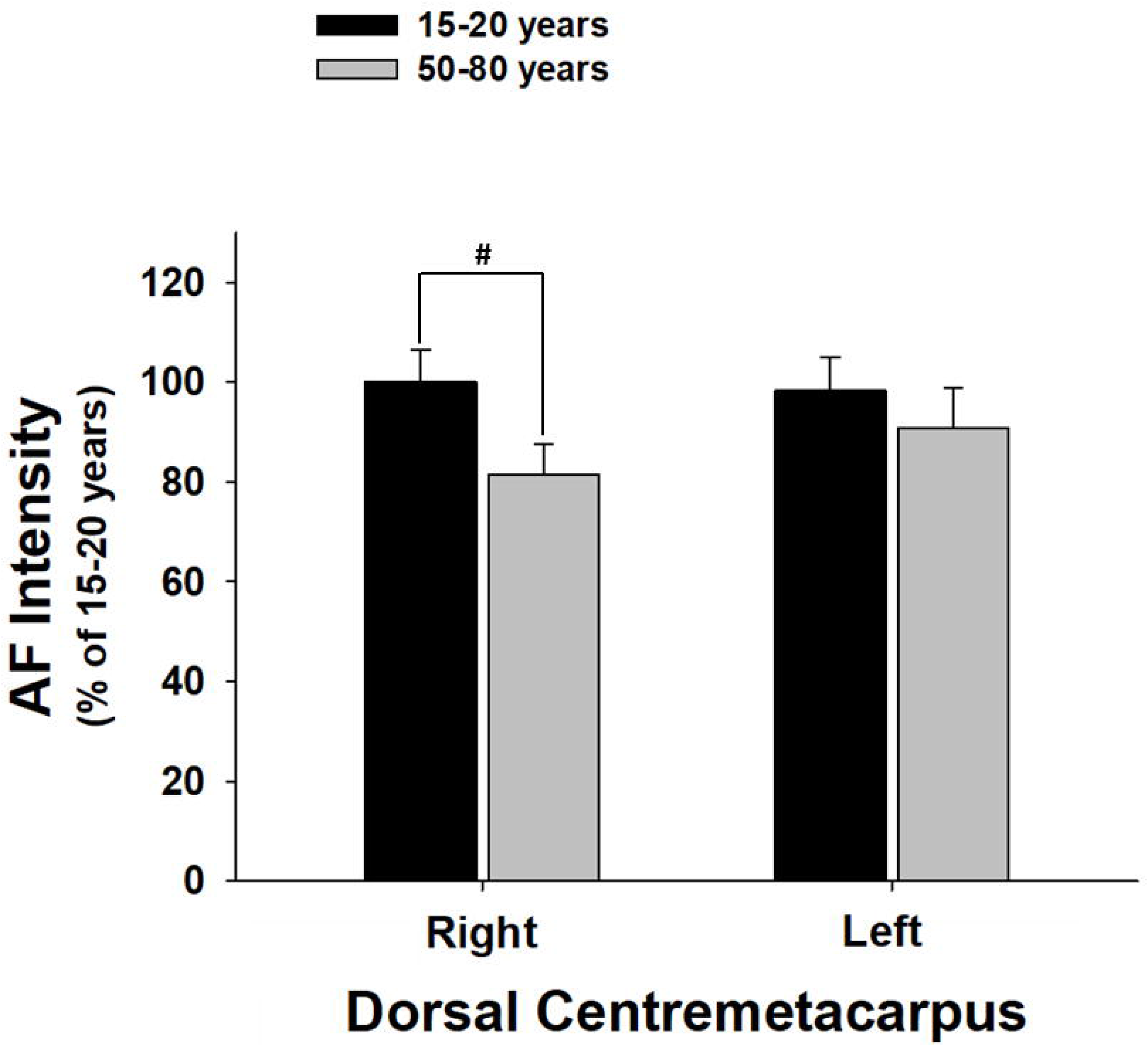
The green AF intensity of the age group between 15 – 20 years of old is significantly higher than that of the age group between 50 – 80 years of old at right Dorsal Centremetacarpus. The number of subjects in the age group between 15 – 20 years of old and the age group between 50 – 80 years of old is 50 and 73, respectively. *, *p* < 0.05.

**Fig. 3.**
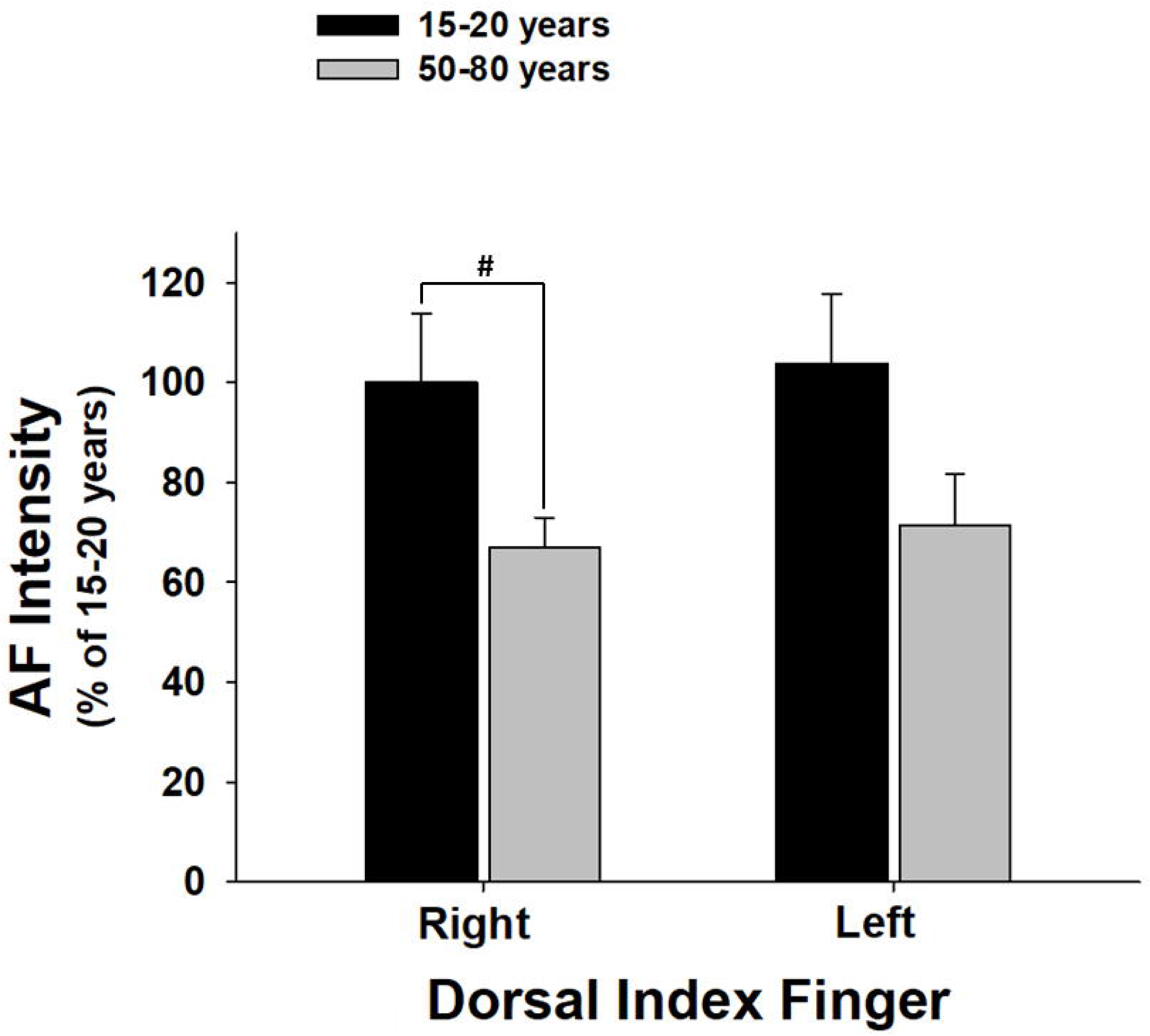
The green AF intensity of the age group between 15 – 20 years of old is significantly higher than that of the age group between 50 – 80 years of old at right Dorsal Index Finger. The number of subjects in the age group between 15 – 20 years of old and the age group between 50 – 80 years of old is 50 and 73, respectively. *, *p* < 0.05.

### 2. The green AF intensity is negatively correlated with the age for certain age groups at three regions examined

For the green AF intensity of the age groups of 15 – 20 years of old, 60 – 70 years of old, 70 – 80 years of old, and 81 – 85 years of old, the green AF intensity is negatively correlated with the age at both right and left Dorsal Index Fingers (Fig. 4) and left Centremetacarpus (Fig. 5). In contrast, no correlation among the green AF intensity of these age groups was found at other regions examined, including Ventroforefinger (Supplemental Fig. 5), Dorsal Centremetacarpus (Supplemental Fig. 6), Ventriantebrachium (Supplemental Fig. 7), Dorsal Antebrachium (Supplemental Fig. 8), and Index Fingernails (Supplemental Fig. 9).

**Fig. 4.**
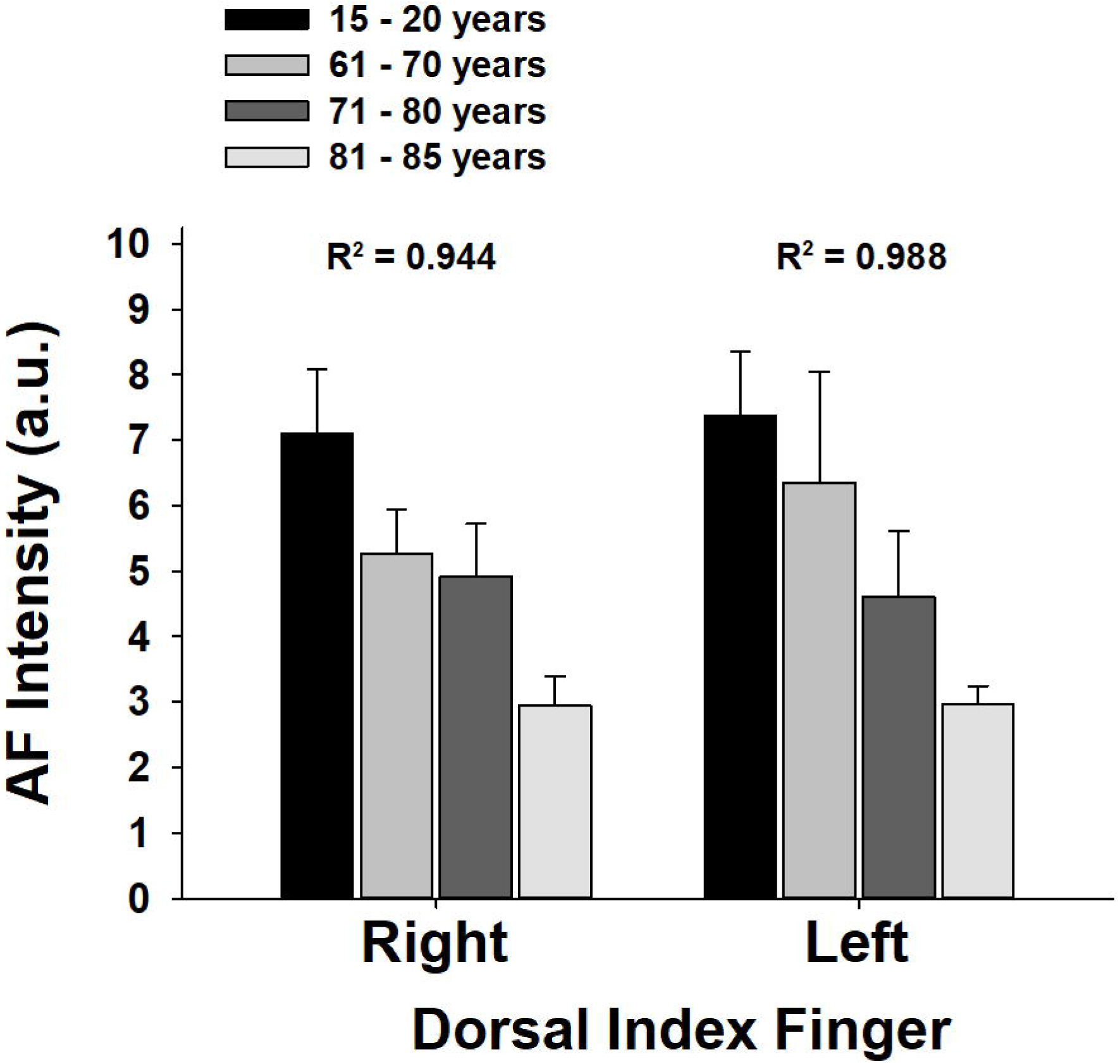
For the green AF intensity of the age groups of 15 – 20 years of old, 60 – 70 years of old, 70 – 80 years of old, and 81 – 85 years of old, the green AF intensity is negatively correlated with the age at both right and left Dorsal Index Fingers. The number of subjects in the age group of 15 – 20 years of old, 60 – 70 years of old, 70 – 80 years of old, and 81 – 85 years of old is 50, 17, 28, 20, and 8, respectively.

**Fig. 5.**
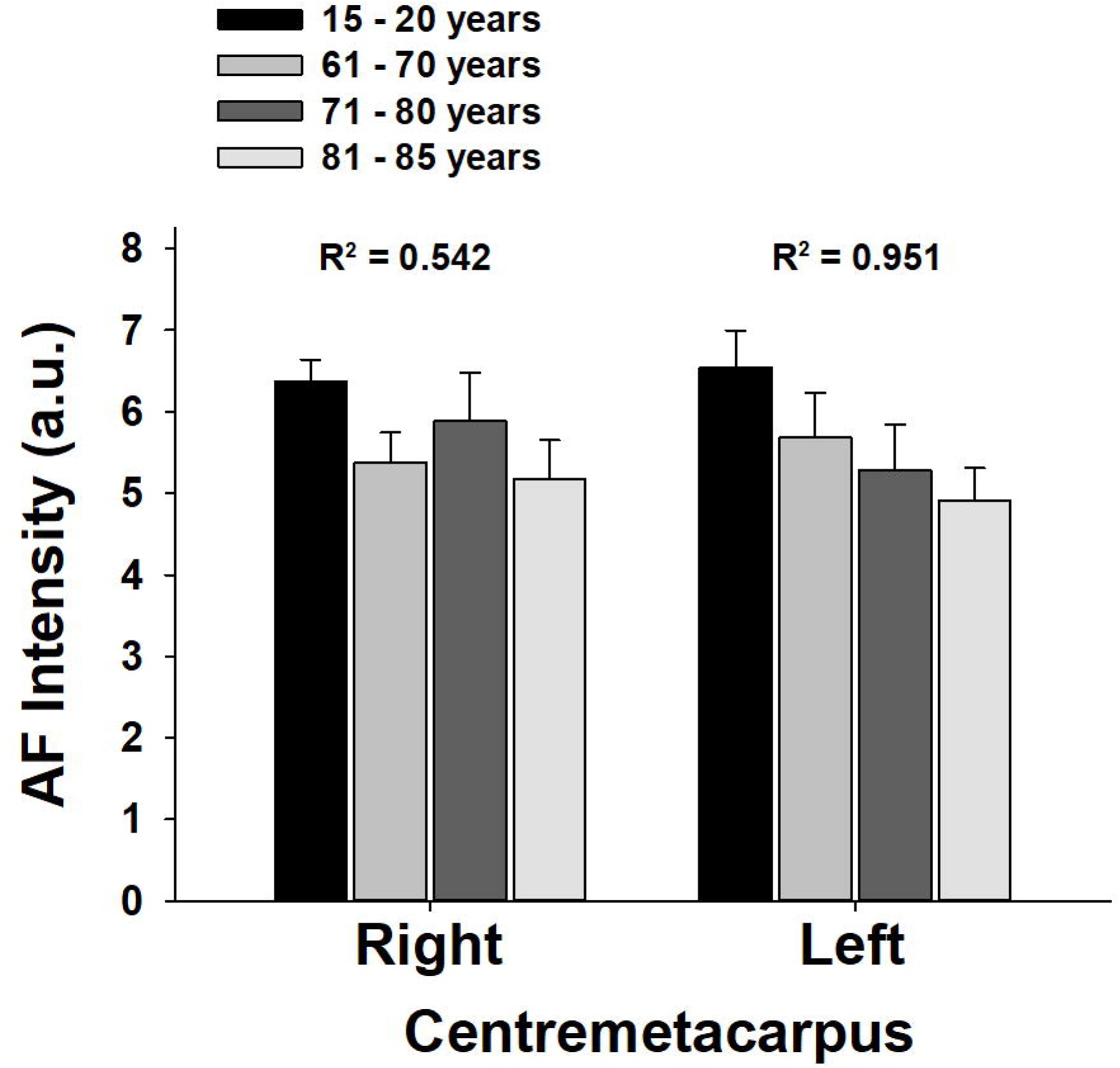
For the green AF intensity of the age groups of 15 – 20 years of old, 60 – 70 years of old, 70 – 80 years of old, and 81 – 85 years of old, the green AF intensity is negatively correlated with the age at left Centremetacarpus. The number of subjects in the age group of 15 – 20 years of old, 60 – 70 years of old, 70 – 80 years of old, and 81 – 85 years of old is 50, 17, 28, 20, and 8, respectively.

## Discussion

The major findings of our current study include: First, the green AF intensity of the age group between 15 - 20 years of old is significantly higher than that of the age group between 50 - 80 years of old at both right and left Centremetacarpus, right Dorsal Centremetacarpus, and right Dorsal Index Finger. Second, for the green AF intensity of the age groups of 15 - 20 years of old, 60 - 70 years of old, 70 - 80 years of old, and 81 - 85 years of old, the green AF intensity is negatively correlated with the age at both right and left Dorsal Index Fingers and left Centremetacarpus. Collectively, our study has provided first evidence showing the age dependence of the AF intensity of human’s skin, which has established essential basis for the studies that determine the diagnostic value of the green AF for age-independent diseases.

Our previous studies have suggested that each of the five major diseases, including AIS (3), MI (17), stable coronary artery disease (17), Parkinson’s disease (4) and lung cancer (10), has its unique ‘Pattern of AF’. In particular, we have found that each of the major diseases has its unique pattern of the increases in the green AF intensity at certain regions of the skin. It is noteworthy that all of these major diseases are age-related diseases. The age of all of the subjects, including healthy controls and the patients of the major diseases, in these studies ranges from 50 – 80 years of old. While all our studies on the ‘Pattern of AF’ have used ‘age-matched’ controls, it has remained unknown if the AF intensity is age-dependent in the healthy controls. Our current study has provided first direct evidence that at least at certain regions of the skin examined, the AF intensity shows age-dependence in the healthy controls. This finding has established guideline for the future studies on the AF-based diagnosis of diseases: The AF intensity of the patients of certain age group should be compared statistically only with the healthy controls of the same age group.

A key scientific question for our current finding is: What is the mechanism underlying the age dependence of the AF intensity ? Our previous studies have suggested that oxidative stress-induced keratin 1 proteolysis is a major cause of the increased green AF intensity of the skin and fingernails of the patients. Therefore, two factors may determine the AF intensity: The first is the levels of oxidative stress in the blood, and the second is the keratin 1 levels in the epidermis. It has been reported that chronic UV radiation – a key cause of skin aging (12) – on the skin can induce a decrease in the keratin 1 level of the skin (14). In contrast, it is established that there is an age-dependent increase in oxidative stress in human body (15). Therefore, we propose that the age-dependent decrease in the keratin 1 levels may play a key role in the age dependence of the AF intensity as observed in our current study.

## Acknowledgment

The authors would like to acknowledge the financial support by a Major Special Program Grant of Shanghai Municipality (Grant # 2017SHZDZX01) (to W.Y. and X.C.), a Major Research Grant from the Scientific Committee of Shanghai Municipality #16JC1400500 and #16JC1400502 (to W.Y. and X.C.), and National Key Research and Development Program Grants of China #2017YFC0907002 and 2017YFC0907501 (to X.C.).

## Legends of Supplemental Figures

**Supplemental Figure 1**. No difference in the green AF intensity at Ventroforefinger between the age group between 15 – 20 years of old and the age group between 50 – 80 years was observed. The number of subjects in the age group between 15 – 20 years of old and the age group between 50 – 80 years of old is 50 and 73, respectively.

**Supplemental Figure 2**. No difference in green AF intensity at Ventriantebrachium between the age group between 15 – 20 years of old and the age group between 50 – 80 years was observed. The number of subjects in the age group between 15 – 20 years of old and the age group between 50 – 80 years of old is 50 and 73, respectively.

**Supplemental Figure 3**. No difference in green AF intensity at Dorsal Antebrachium between the age group between 15 – 20 years of old and the age group between 50 – 80 years was observed. The number of subjects in the age group between 15 – 20 years of old and the age group between 50 – 80 years of old is 50 and 73, respectively.

**Supplemental Figure 4**. No difference in green AF intensity at Index Fingernails between the age group between 15 – 20 years of old and the age group between 50 – 80 years was observed. The number of subjects in the age group between 15 – 20 years of old and the age group between 50 – 80 years of old is 50 and 73, respectively.

**Supplemental Figure 5**. For the green AF intensity of the age groups of 15 – 20 years of old, 60 – 70 years of old, 70 – 80 years of old, and 81 – 85 years of old, no correlation among the green AF intensity of these age groups was found at Ventroforefinger. The number of subjects in the age group of 15 – 20 years of old, 60 – 70 years of old, 70 – 80 years of old, and 81 – 85 years of old is 50, 17, 28, 20, and 8, respectively.

**Supplemental Figure 6**. For the green AF intensity of the age groups of 15 – 20 years of old, 60 – 70 years of old, 70 – 80 years of old, and 81 – 85 years of old, no correlation among the green AF intensity of these age groups was found at Dorsal Centremetacarpus. The number of subjects in the age group of 15 – 20 years of old, 60 – 70 years of old, 70 – 80 years of old, and 81 – 85 years of old is 50, 17, 28, 20, and 8, respectively.

**Supplemental Figure 7**. For the green AF intensity of the age groups of 15 – 20 years of old, 60 – 70 years of old, 70 – 80 years of old, and 81 – 85 years of old, no correlation among the green AF intensity of these age groups was found at Ventriantebrachium. The number of subjects in the age group of 15 – 20 years of old, 60 – 70 years of old, 70 – 80 years of old, and 81 – 85 years of old is 50, 17, 28, 20, and 8, respectively.

**Supplemental Figure 8**. For the green AF intensity of the age groups of 15 – 20 years of old, 60 – 70 years of old, 70 – 80 years of old, and 81 – 85 years of old, no correlation among the green AF intensity of these age groups was found at Dorsal Antebrachium. The number of subjects in the age group of 15 – 20 years of old, 60 – 70 years of old, 70 – 80 years of old, and 81 – 85 years of old is 50, 17, 28, 20, and 8, respectively.

**Supplemental Figure 9**. For the green AF intensity of the age groups of 15 – 20 years of old, 60 – 70 years of old, 70 – 80 years of old, and 81 – 85 years of old, no correlation among the green AF intensity of these age groups was found at Index Fingernails. The number of subjects in the age group of 15 – 20 years of old, 60 – 70 years of old, 70 – 80 years of old, and 81 – 85 years of old is 50, 17, 28, 20, and 8, respectively.

